# Parsing propagule pressure: Number, not size, of introductions drives colonization success in a novel environment

**DOI:** 10.1101/108324

**Authors:** Michael J Koontz, Meagan F Oldfather, Brett A Melbourne, Ruth A Hufbauer

**Affiliations:** Graduate Group in Ecology, University of California, Davis, CA 95616; Graduate Degree Program in Ecology, Colorado State University, Fort Collins, CO 80523; Department of Bioagricultural Science and Pest Management, Colorado State University, Fort Collins, CO 80523; Department of Integrative Biology, University of California, Berkeley, CA 94720; Department of Ecology and Evolutionary Biology, University of Colorado, Boulder, CO 80309

**Keywords:** invasion, reintroduction, propagule pressure, biocontrol, conservation, microcosm, population dynamics, stochasticity, simulation

## Abstract

Predicting whether individuals will colonize a novel habitat is of fundamental ecological interest and is crucial to both conservation efforts and invasive species management. A consistently supported predictor of colonization success is the number of individuals introduced, also called propagule pressure. Propagule pressure increases with the number of introductions and the number of individuals per introduction (the size of the introduction), but it is unresolved which process is a stronger driver of colonization success. Furthermore their relative importance may depend upon the environment, with multiple introductions potentially enhancing colonization of fluctuating environments. To evaluate the relative importance of the number and size of introductions and its dependence upon environmental variability, we paired demographic simulations with a microcosm experiment. Using *Tribolium* flour beetles as a model system, we introduced a fixed number of individuals into replicated novel habitats of stable or fluctuating quality, varying the number of introductions through time and size of each introduction. We evaluated establishment probability and the size of extant populations after 7 generations. In the simulations and microcosms, we found that establishment probability increased with more, smaller introductions, but was not affected by biologically realistic fluctuations in environmental quality. Population size was not significantly affected by environmental variability in the simulations, but populations in the microcosms grew larger in a stable environment, especially with more introduction events. In general, the microcosm experiment yielded higher establishment probability and larger populations than the demographic simulations. We suggest that genetic mechanisms likely underlie these differences and thus deserve more attention in efforts to parse propagule pressure. Our results highlight the importance of preventing further introductions of undesirable species to invaded sites, and suggest conservation efforts should focus on increasing the number of introductions or re-introductions of desirable species rather than increasing the size of those introduction events.

## INTRODUCTION

Colonization is the ecologically fundamental process of population establishment in an unoccupied site, and it underlies the past, present, and future distributions of species. Colonization occurs naturally, but is increasingly prevalent due to anthropogenic influences (Sakai et al. 2001, Cassey et al. 2005, Ricciardi 2007). Incipient populations often face environments that are entirely novel, which is especially likely in the case of anthropogenic colonization (Cassey et al. 2005, Ricciardi 2007). Regardless of whether colonization events to novel habitats are natural (e.g., range expansion) or human-mediated (e.g., biological invasions, reintroductions of rare species, release of biological control agents), their successes or failures have significant implications for natural resource managers and society (Mack et al. 2000).

Most introductions to novel habitats fail, and colonization success can be difficult to predict (Lockwood et al. 2005, Zenni and Nuñez 2013). Incipient populations are commonly small, and face threats from environmental, demographic, and genetic stochasticity (Lande 1988, 1993, Fauvergue et al. 2012). Furthermore, the success of any given population can be idiosyncratic with respect to taxonomy and geography (Lodge 1993, Lockwood et al. 2005). Thus, it is crucial to understand more general features of the colonization process beyond the particular invading organism or the particular invaded environment (Lockwood et al. 2005). Propagule pressure is one such general feature that is a consistent predictor of colonization success in novel habitats (Lockwood et al. 2005, Colautti et al. 2006, Simberloff 2009, Jeschke 2014).

Propagule pressure is the total number of potentially reproductive individuals (e.g., adults, eggs, seeds, vegetative material) introduced to an area (Novak 2007). It is often described in this broad sense, which belies its complexity. Two important components of propagule pressure are the number of introduction events— sometimes termed propagule number, and the number of individuals per introduction event— sometimes termed propagule size (*sensu* Fauvergue et al. 2012). Here, we use “introduction regime” to refer to different combinations of the number of introduction events and the number of individuals introduced per event. The same total propagule pressure, *N*, is realized by different introduction regimes depending on how those *N* individuals are distributed in time or space (Haccou and Iwasa 1996). An introduction regime of *N* individuals lies on a continuum bounded by maximizing the number of individuals introduced per event (all *N* individuals introduced in one event to the same location) and maximizing the number of introduction events (one individual introduced in each of *N* sequential events through time or to each of *N* unique locations).

Colonization success increases with total propagule pressure, but it is unclear whether the correlation is driven by the number of individuals introduced per event or the number of introduction events (Lockwood et al. 2005, Colautti et al. 2006, Simberloff 2009). Historical data suggest that multiple introductions can facilitate population establishment, but conclusions from these studies are limited by the inability to control for the total number of individuals introduced (Hopper and Roush 1993, Blackburn and Duncan 2001, Grevstad et al. 2011, Fauvergue et al. 2012). Models that hold the total propagule pressure constant agree that multiple, small introductions distributed across space will lead to greater establishment probability compared to a single, large introduction when Allee effects are weak (Haccou and Iwasa 1996, Grevstad 1999, Schreiber and Lloyd-Smith 2009). However, both modeling and empirical approaches have generated conflicting views on how an introduction regime affects colonization success when introductions are distributed through time, a situation in which individuals from later introductions interact both demographically and genetically with individuals from previous introductions.

There is evidence from both models and experiments that colonization success can increase with more, smaller introduction events through time. Branching process models show that, in the long run, several small introductions will be more likely to successfully establish a population than a single large introduction (Haccou and Iwasa 1996, Haccou and Vatutin 2003). This finding was corroborated by simulations with no Allee effects (Drolet and Locke 2016). In a *Daphnia* microcosm experiment, increasing introduction frequency (proportional to the number of introductions) positively affected population growth, but the number of individuals per introduction had no detectable effect (Drake et al. 2005). Establishment probability was only affected by total propagule pressure, and did not increase with increasing introduction frequency (Drake et al. 2005). However, Drake et al. (2005) did not continue scheduled introductions if a population went extinct, denying those populations one of the main benefits of repeated introductions and creating variability in the total propagule pressure. In a more recent experiment, several small introductions through time led to a 65% increase in abundance of successfully colonizing invasive Pacific Oyster (*Crassostrea gigas*) compared to a single large introduction (Hedge et al. 2012).

There is contrasting evidence from both models and experiments that colonization success increases with fewer, larger introduction events through time. In simulations of bird invasions, a single large introduction event always led to the greatest establishment probability (Cassey et al. 2014). When Drolet and Locke (2016) included Allee effects in their simulations, their previously observed pattern reversed and colonization success was instead more likely with fewer, larger introductions than with more, smaller introductions. In a field experiment with the psyllid biocontrol agent, *Arytainilla spartiophila*, the number of individuals per introduction event was a better predictor of establishment success than the number of introduction events (Memmott et al. 2005). In this case however, the introduction regimes with the most individuals per event also had the highest total propagule pressure (Memmott et al. 2005). In an experiment that controlled total propagule pressure, a single, large introduction of the non-native mysid, *Hemimysis anomala*, led to larger populations and greater survival probabilities compared to several, small introductions through time (Sinclair and Arnott 2016).

Environmental stochasticity in the recipient environment may also affect which introduction regime is optimal for colonization. Branching process models show that a more variable environment reduces the probability of population establishment for all introduction regimes (Haccou and Iwasa 1996, Haccou and Vatutin 2003), and simulations of introductions distributed in space suggest that greater environmental variability will magnify the benefit of multiple introductions (Grevstad 1999). However, simulations of introductions through time by Cassey et al. (2014) did not support either of these outcomes— a single, large introduction was most likely to establish a population even with extreme levels of environmental stochasticity.

Thus, modeling and empirical approaches have not resolved how different introduction regimes with a fixed total propagule pressure will affect colonization success when introductions are distributed through time. Further, there has been no experimental test of whether variability in the recipient environment interacts with the introduction regime to affect colonization. We paired demographic simulations with a laboratory microcosm experiment using the red flour beetle, *Tribolium castaneum*, to reconcile conflicts in the literature and test how different introduction regimes implemented through time affect colonization success in novel habitats. We explicitly manipulated whether the novel habitat was stable or randomly fluctuating in quality to assess how the success of different introduction regimes may depend upon variability in the recipient environment. With the total number of individuals introduced held constant at 20, we varied the size and number of introduction events used to distribute those individuals in four different introduction regimes. We evaluated establishment probability and population size over 7 discrete generations to ask: 1) does colonization success increase with more introduction events or with more individuals per introduction event?, and 2) does the effect of the introduction regime on colonization success depend on whether the recipient novel environment is stable or fluctuating through time?

## METHODS

### General Framework

In simulations and a microcosm experiment, we evaluated the outcome of introducing 20 total individuals to one of two environmental contexts (a stable or fluctuating novel environment), varying the number of introduction events used to distribute those individuals through time. This total propagule pressure was low enough to allow some population extinction within an observable timeframe, but high enough to be representative of documented introductions in the literature (Simberloff 1989, Simberloff 2009; Grevstad 1999; Berggren 2001; Taylor et al. 2005; Drake et al. 2005). The introduction regimes were: 20 individuals introduced in the first generation, 10 individuals introduced in each of the first 2 generations, 5 individuals introduced in each of the first 4 generations, and 4 individuals introduced in each of the first 5 generations. To create the fluctuating environment, we imposed a magnitude of variability corresponding to environmental stochasticity in nature, which leads to frequent, mild to moderate perturbations in population growth rate due to external forces (Lande 1993). Populations were tracked for seven generations following the initial introduction.

Establishment probability and the size of established populations were used as measures of colonization success. Populations were deemed ‘established’ for any time step in which they were extant and population size was noted for all extant populations in every time step. Establishment probability and mean population size were assessed in generation 7.

### Study System

Our simulations and microcosm experiment use *Tribolium castaneum* (red flour beetles) to model the life history of organisms with discrete, non-overlapping generations (e.g., annual insects and fishes) following Melbourne and Hastings (2008). Individual simulated colonists were randomly sourced from a randomly mating, infinitely large population. Individual experimental colonists came from a thoroughly mixed source population maintained at 800 individuals in each of the 4 temporal blocks over which the experiment was replicated. To obtain colonists, beetles from the source population were mixed freely on a plate, selected from many sections of the plate, and introduced to subsets of the experiment populations in a random order (about 24 subsets per block). All migrant females were assumed to arrive mated from the source population. The strong maternal effects exhibited by *Tribolium* flour beetles were reduced by rearing individuals from the source population on novel growth medium for 1 generation prior to using them as colonists (Van Allen and Rudolf 2013, Hufbauer et al. 2015).

### Simulations

Population dynamics were simulated with a Negative Binomial-Binomial gamma (NBBg) Ricker model (Melbourne and Hastings 2008) with an additional mating function of the form:

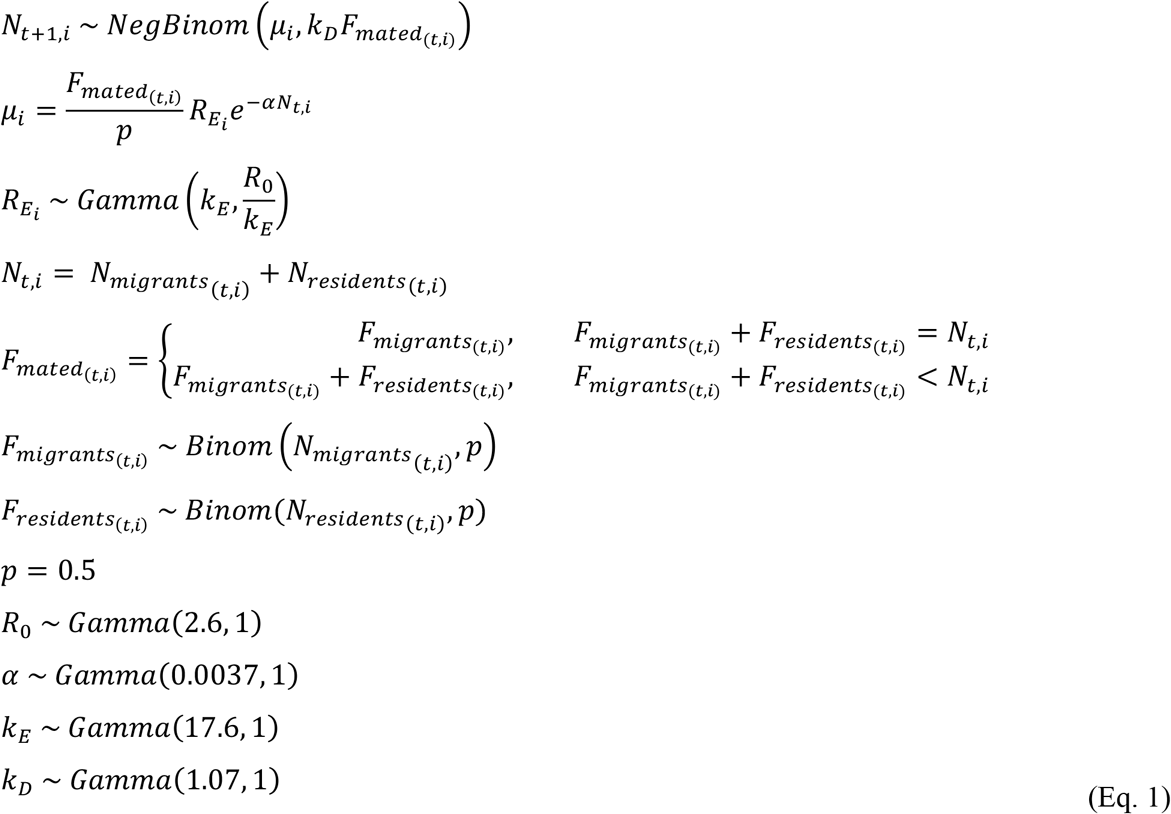

Where *N*_*t+1,i*_ is the size of population *i* at time *t*+*1*, *k*_*D*_ represents stochasticity due to demographic heterogeneity, *μ*_*i*_ is the expectation of the size of population *i* in the next time step, *F*_*mated(t,i)*_ is the latent number of mated females in population *i* at time *t*, *p* is the probability of an individual being female, *R*_*Ei*_ is the latent density-independent population growth rate for environment *E*, *α* is the egg cannibalism rate, *N*_*t,i*_ is the total size of population *i* at time *t*, *k*_*E*_ represents environmental stochasticity, *R*_*0*_ is the density-independent population growth rate for the average environment, *N*_*migrants(t,i)*_ is the total number of migrants to population *i* at time *t*, and *N*_*residents(t,i)*_ is the total number of residents in population *i* at time *t*.

We imposed a mating function such that a population would deterministically go extinct if it comprised only non-migrant females. In cases with an all female population and a mixture of residents and migrants, we only included the number of migrant females in the density-dependent effect of the demographic heterogeneity term *k*_*D*_\**F*_*mated(t,i)*_ because this effect manifests via the number of eggs laid by females and only migrant females, being pre-mated, would lay eggs. Weakly regularizing gamma priors were taken from Melbourne and Hastings (2008). The expected equilibrium population size for the model is:

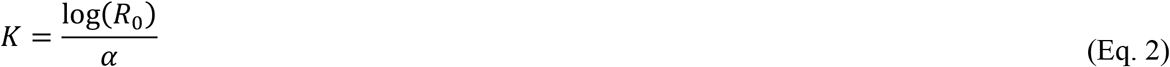

For each combination of the 4 introduction regimes and 2 recipient environment types (described above), we simulated 500,000 replicate populations for 7 generations. All simulations were performed in R (R Core Team 2016). This model captures the stochastic and deterministic ecological dynamics of the *Tribolium* system well, but does not include evolutionary processes (Melbourne and Hastings 2008, Hufbauer et al. 2015).

To parameterize the model, we censused 125 *Tribolium* populations one generation after establishing them at various, known densities (between 5 and 200 individuals) on the novel growth medium following the rearing procedure described in “Microcosm Experiment” below. We fit the hierarchical model (Eq. 1) to these data in a Bayesian framework to generate posterior distributions for each of the parameters using the nimble package in R (de Valpine et al. 2016). We used a Metropolis-Hastings random walk sampler with 3 chains having 50,000 samples each (including 10,000 samples that were removed for burn in). The chains converged (multivariate 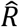 for the 4 key parameters =1.01) and produced a sufficient number of effective samples (*R*_*0*_: 757, *α*: 548, *k*_*E*_: 4103, *k*_*D*_: 1703).

A single novel growth medium mixture was used for these 125 populations (mixture 5, Supplementary Table 1), so we treated the estimated environmental stochasticity parameter, *k*_*E*_, as the expectation for a stable environment. We parameterized a fluctuating environment model by increasing the standard deviation of the density-independent per capita population growth rate by approximately 10% for any given population at any given time step. This was accomplished by multiplying all of the samples from the posterior distribution of the environmental stochasticity parameter, *k*_*E*_, by 100/121 (derived by moment matching for the gamma distribution). By using random sets of parameters drawn from the samples of the posterior distributions, we were able to propagate the model uncertainty through the simulations and combine it with the deterministic and stochastic dynamics represented by the model itself.

### Microcosm Experiment

We founded 842 *Tribolium* populations with different introduction regimes (20 individuals in the first generation, 10 individuals in the first 2 generations, 5 individuals in the first 4 generations, or 4 individuals in the first 5 generations) and environments (stable or fluctuating) with between 96 and 120 replicate populations per treatment combination (Supplemental Table 3). Each population was reared in a 4cm × 4cm × 6cm plastic box (AMAC Plastic Products) with 2 tablespoons (approximately 15 g) of freshly prepared growth medium that had been humidified for at least 24 hours. The growth medium used for the source population (the natal environment of the colonists) comprised 95% wheat flour (Pillsbury Co. or Gold Medal Products Co.) and 5% brewers' yeast (Sensient Flavors). Colonists were introduced into a novel growth medium comprising a small percentage of natal medium mixed with corn flour (Bob’s Red Mill). All populations were reared in one of two dark incubators at 31° and approximately 70% relative humidity (standard conditions) and were haphazardly rotated between incubators weekly.

For each population in each generation, a known number of adults laid eggs for 24 hours in fresh medium, and were then removed. Offspring were given 35 days to develop, and adults were then censused. Censused adults laid eggs on freshly prepared growth medium for 24 hours, completing their laboratory life cycle. We estimated the maximum observation error during census to be 4.6% (median: 0%, mean: 0.26%) with no detectable difference across observers or populations with different sizes.

Each replicate population experienced a novel environment that was either stable or randomly fluctuating through time. The same novel growth medium mixture containing 99.05% novel corn flour and 0.95% natal medium (mixture 5, Supplementary Table 1) was used for the stable environment for the duration of the experiment, which preliminary results indicated would yield a population growth rate of λ = 1.2 compared to a mean population growth rate of 3.36 on 100% natal medium. To create the fluctuating environment, we randomly selected a novel growth medium mixture from one of 9 possible media mixtures for each population in each generation. Each population in the fluctuating environment therefore experienced a unique series of environmental conditions. The 9 possible media mixtures represented a gradient of corn flour to natal medium ratios, and were designed to yield expected population growth rates between 0.88 and 1.33 (Supplementary Table 1). We chose this range to mimic environmental stochasticity measured in nature (Sæther and Engen 2002) while remaining within the bounds of biologically realistic population growth (λ between 0.5 and 1.5) (Morris et al. 2008). The random sequence of growth media was constrained such that the expected geometric mean population growth rate for each population resembled expected growth of populations in the stable environment (*λ_expected_* = 1.2±0.05).

We estimated the amount of environmental stochasticity that we achieved in the fluctuating environment as the difference in mean total stochasticity between populations in the fluctuating and stable environments (Sæther and Engen 2002). We assumed that total stochasticity was a combination of demographic and environmental stochasticity for populations in the fluctuating environment, and that demographic stochasticity was the sole contributor to total stochasticity for populations in the stable environment. Total stochasticity (demographic plus environmental) of each population that did not experience extinction (n=667) was calculated as the variance of the natural logarithms of its population growth rates through 7 generations:

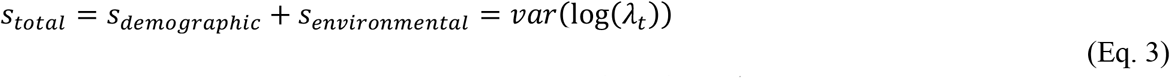
where, for a particular population, *S*_*total*_ is its total stochasticity, *S*_*demographic*_ is its demographic stochasticity, *S*_*environmental*_ is its environmental stochasticity (assumed to be 0 for populations in the stable environment), and *λ*_*t*_ is its per capita population growth rate between generation *t-1* and generation *t* (*t=1, 2, …, 7*). We only calculated total stochasticity for populations that did not experience any extinction in order to capture the full temporal extent of environmental fluctuations and because extinctions would have an infinite effect on this measure of stochasticity.

### Statistical analyses

We evaluated how our environment treatment affected variability in population growth rate (total stochasticity from Eq. 3) using a linear mixed effects model with environment (stable or fluctuating) as a fixed effect and block as a random intercept effect.

We used a mixed effects logistic regression with a logit link to predict the binary response of establishment, and a mixed effects Poisson regression with a log link to analyze population size. In both models, introduction regime, environment treatment, and their interaction were treated as fixed effects, and block was treated as a random intercept effect.

We assessed the effect of temporary extinctions on the establishment probability and mean population size by fitting the generalized linear mixed effects models described above to data from the multiple introduction regimes (i.e. not the 20×1 regime) and with additional predictor variables. To assess the effect of the presence of a temporary extinction, we included an additional Boolean predictor for whether a population went temporarily extinct or not. To assess the role of the total propagule pressure, we included a numeric predictor representing the number of beetles introduced after the latest temporary extinction.

Group-level significance of fixed effects was tested using likelihood ratio tests on nested models, and least-squares contrasts were used to compare levels of the fixed effects. All statistical analyses were performed in R, version 3.3.2 (R Core Team 2016). Generalized linear mixed models were fit using the lme4 package, version 1.1-12 (Bates et al. 2015) and pairwise comparisons were made using the lsmeans package, version 2.25 (Lenth 2016).

### Data availability

The raw experiment data, simulated population trajectories, R code for the simulations, R code fitting the NBBg model, samples from the posterior distributions of the NBBg parameters, and R code fitting the mixed effects models are available as supplemental material, on Figshare (https://doi.org/10.6084/m9.figshare.4648865.v1), and on GitHub (https://github.com/mikoontz/ppp-establishment).

### Note on egg contamination

Laboratory procedures after generation 3 resulted in occasional egg contamination between replicate populations of the same treatment. Some populations of 1 individual persisted when they should have deterministically gone extinct, so we manually edited those population trajectories such that they went extinct in the generation after having only 1 individual.

## RESULTS

### Simulations

Summary statistics for the posterior distributions of the four NBBg model parameters are given in Supplementary Table 2.

Our simulations showed introduction regimes with more introduction events were more likely establish a population by the seventh generation (Figure 1). Simulated introductions into a stochastically fluctuating environment resulted in slightly lower population establishment for all introduction regimes (difference of ~1%), but did not favor any particular regime (not shown). The mean population sizes for each introduction regime/environment combination were approximately equal in simulations. Mean population sizes only incorporated extant populations, so they were slightly larger than the expectation for the equilibrium population size (Figure 2).

**Figure 1.**
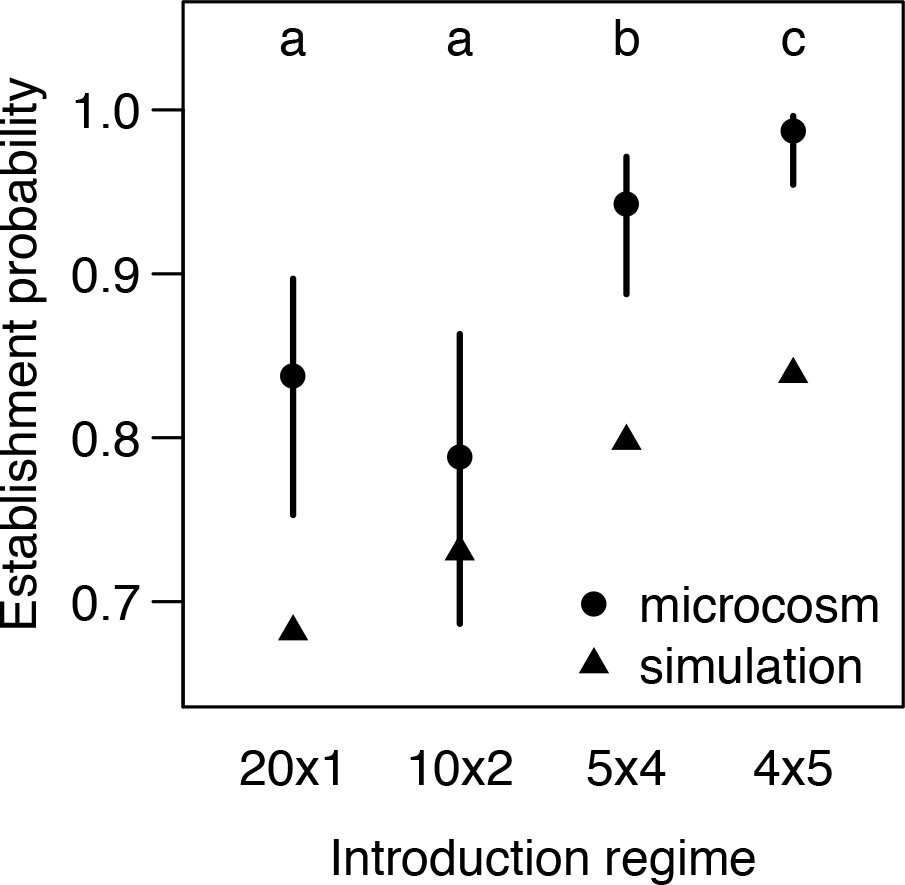
Establishment probabilities for each introduction regime assessed at generation 7. Triangles represent results from simulations. Dot-whiskers represent estimates and 95% confidence intervals from the mixed effects logistic regression model fit to data from the microcosm experiment. Different letters over dot-whiskers represent introduction regimes with significantly different establishment probabilities in the microcosm experiment.

**Figure 2.**
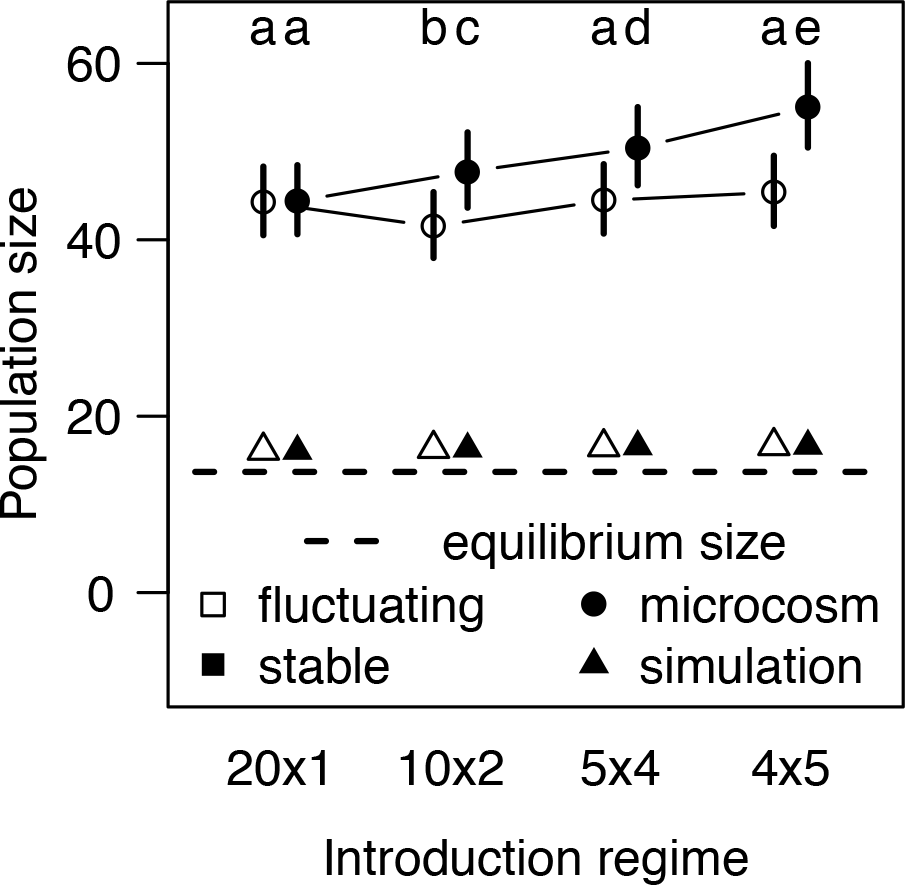
Mean sizes of extant populations for each introduction regime/environment combination in generation 7. Triangles represent results from simulations. Dot-whiskers represent estimates and 95% confidence intervals from the mixed effects Poisson regression model fit to data from the microcosm experiment. Different letters over dot-whiskers represent introduction regimes with significantly different mean population sizes. The dashed line represents the theoretical equilibrium population size derived using Eq. 2, which includes extinction and is therefore lower than the mean population size from the simulations.

### Microcosm experiment

Mean environmental stochasticity of populations in the fluctuating environment was 0.052 (95% CI = 0.0073 to 0.0966; p = 0.023). This value is near the median value of 0.055 measured in nature by Sæther and Engen (2002) in a metaanalysis of 35 avian populations, indicating that we achieved biologically realistic fluctuations in population growth rate.

We found no evidence that the probability of establishment was affected by a main effect of environment (χ^2^=0.72, df=1, p=0.40), nor by an interaction between environment and introduction regime (χ^2^=3.49, df=3, p=0.32). However, there was strong support for an effect of introduction regime on establishment probability (χ^2^=59.76, df=3, p<0.0001). Pairwise comparisons of the different introduction regimes averaged across the environment treatments revealed that the 4×5 regime was the most likely to establish populations, with a probability of about 0.98, whereas the 20×1 and 10×2 regimes were the least likely to establish populations, with a probability reduced to about 0.8 (Figure 1, Figure 3).

**Figure 3.**
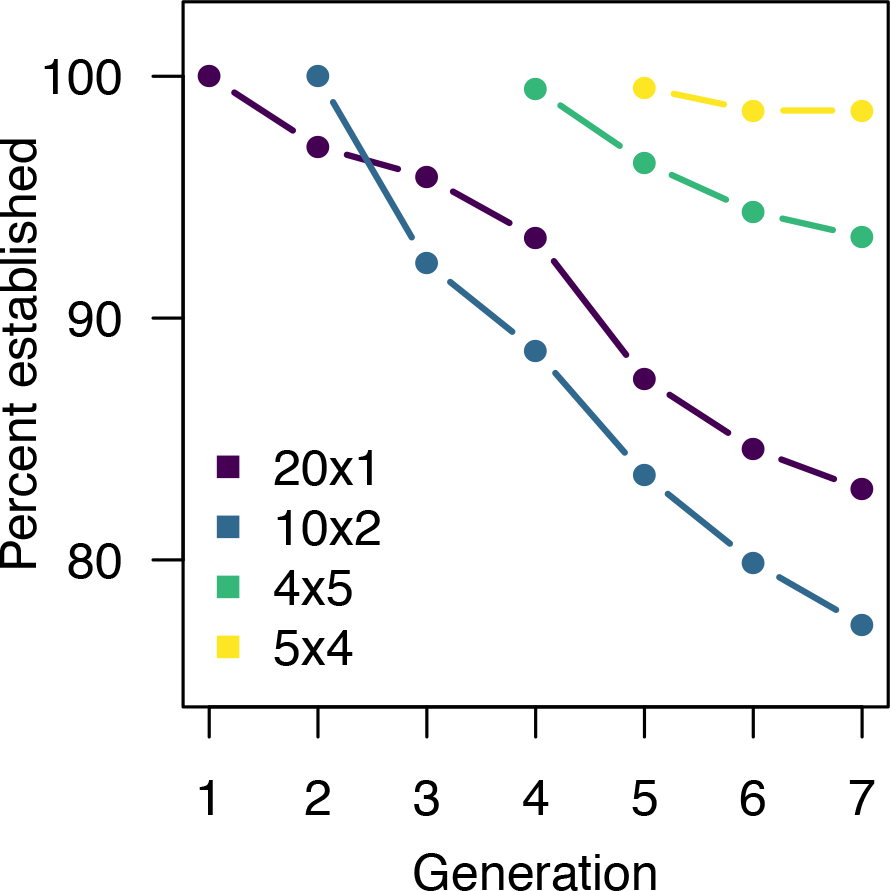
Percent of microcosm populations that were established in each generation for the 4 different introduction regimes. Data are pooled across the two environmental variability treatments. The introduction regimes are as follows: 20×1 = 20 individuals in the first generation, 10×2 = 10 individuals in each of the first two generations, 5×4 = 5 individuals in each of the first 4 generations, and 4×5 = 4 individuals in each of the first 5 generations.

The size of populations that persisted until generation 7 was shaped by significant effects of introduction regime (χ^2^=91.65, df=3, p<0.0001), environment treatment (χ^2^=117.83, df=3, p<0.0001), and their interaction (χ^2^=44.62, df=3, p<0.0001). Populations established via more introduction events were larger when averaged across the environment treatments. Extant populations in the stable environment were larger than those in the fluctuating environment when averaged across the introduction regimes. The interaction manifests as the benefit of a stable environment increases with more, smaller introduction events (Figure 2, Figure 4).

**Figure 4.**
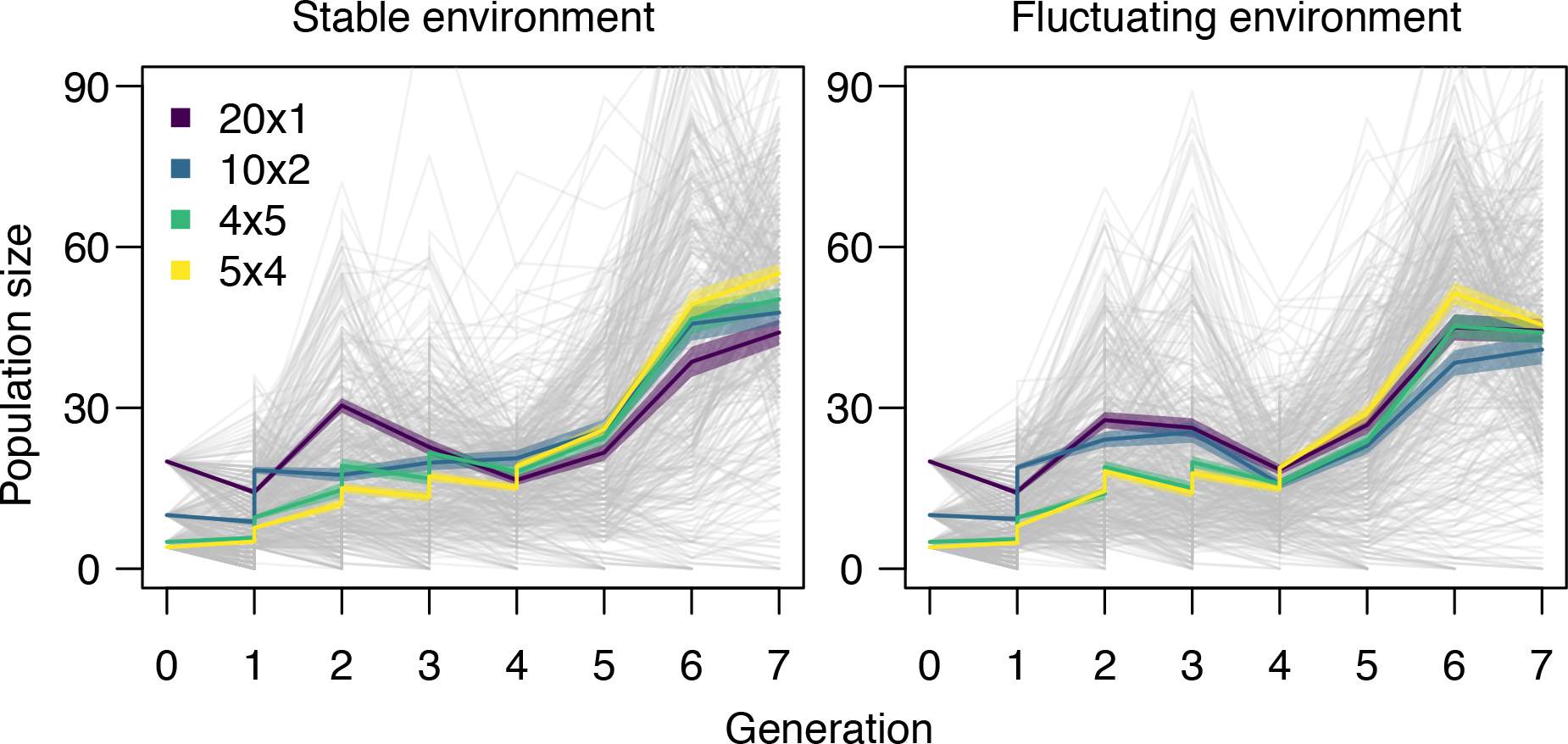
Population trajectories for 842 populations of *Tribolium* in the microcosm experiment. The first panel represents all populations in stable environments (n = 418) and the second panel represents all populations in fluctuating environments (n = 424). Vertical steps in the population size represent additional migrant individuals in introduction regimes with more than 1 introduction. Light grey lines represent individual population trajectories, and colored lines represent the mean size of extant populations for each of the 4 introduction regimes. The introduction regimes are as follows: 20×1 = 20 individuals in the first generation, 10×2 = 10 individuals in each of the first two generations, 5×4 = 5 individuals in each of the first 4 generations, and 4×5 = 4 individuals in each of the first 5 generations. Shading around the colored lines represents the mean size of extant populations plus and minus one standard error.

Extinctions accumulated regularly throughout the experiment period, with 101 out of 842 populations (12.0%) going extinct by generation 7 (Figure 3). The additional introductions that some populations received often restored a population that had temporarily gone extinct. Out of 602 populations that received more than 1 introduction (i.e. not the 20×1 introduction regime), 104 of them (17.3%) temporarily went extinct at least once before being replenished by additional colonizing individuals. Twelve populations were rescued in this way at least twice, and one population was rescued in this way three times.

Temporary extinctions significantly affected colonization success. The presence of a temporary extinction significantly reduced average establishment probability from 92.4% to 82.1% (difference = -1.13 on logit scale, 95% CI = -0.22 to -2.04, χ^2^ = 5.44, df = 1, p = 0.020) and mean population size from 47.8 to 45.4 (difference = -0.052 on log scale, 95% CI = -0.02 to -0.09, χ^2^ = 9.42, df = 1, p = 0.0021). Each additional colonist contributing to a population after the latest temporary extinction significantly increased the mean population size (estimate = 0.005 on the log scale, 95% CI = 6.4e-05 to 0.01, χ^2^ = 3.94, df = 1, p = 0.047).

Results from the simulations are expected to represent the dynamics in the microcosm experiment if the assumption of the model on which the simulations were built holds—that only demographic processes are responsible for the dynamics observed in the experiment. However, establishment probability at generation 7 was equal to or greater than that expected from the simulations (Figure 1) and mean population sizes were much larger in the experiment than in the simulations (Figure 2), together suggesting an influence of non-demographic mechanisms.

## DISCUSSION

We assessed how the number and size of introduction events through time drive colonization success in a novel environment when the total number of individuals introduced to a site is fixed. We considered novel environments that were either stable or randomly fluctuating in quality through time, and evaluated populations through 7 discrete generations. We approached this question in two ways: 1) stochastic simulations of a demographic population dynamics model parameterized with empirical data, and 2) a highly replicated laboratory microcosm experiment. By coupling these approaches, we were able to expand and test the theoretical understanding of how the introduction regime affects colonization in stable and fluctuating environments as well as to develop new avenues for research when observations did not align with predictions. We found that several, small introductions increase colonization success and that demographic processes alone are insufficient to explain the dynamics observed in the experiment.

We found minimal to no effect of a biologically realistic level of environmental stochasticity on establishment probability in demographic simulations or in the microcosm experiment. Our results corroborate those of Cassey et al. (2014) who simulated introductions through time and also found a minimal effect of environmental stochasticity on establishment probability. Cassey et al. (2014) further found a minimal effect of random, infrequent catastrophes and bonanzas, suggesting that increasing temporal environmental variability in the broadest sense (i.e., encompassing environmental stochasticity, random catastrophes, and bonanzas) does not magnify the benefit of several, small introductions through time. A minimal role of environmental stochasticity contrasts with the results of Grevstad (1999) who found with simulations that several, small introductions would produce an especially high establishment probability compared to a single, large introduction in a variable environment. The difference in findings is perhaps due to a difference in the kinds of introduction regimes modeled: Grevstad (1999) simulated multiple introductions in space with strong environmental stochasticity often leading to catastrophic mortality, while we focused on introductions separated in time with less extreme environmental stochasticity leading to moderate fluctuations in growth rates (Lande 1993).

There also did not appear to be a strong effect of environmental stochasticity on population size in the demographic simulations. All treatments in the simulations reached similar mean population size by generation 7, which was slightly higher than the equilibrium size due to our measure only including extant populations. This suggests that strong density dependence in the demographic simulations was the primary influence on population size.

Population growth rate may interact with introduction regime to affect colonization success. Cassey et al. (2014) suggested that their lower simulated mean population growth rates, where *R* was between 1.0 and 1.38, compared to that of Grevstad (1999), where *R* was equal to 2.0, explained why they found that a single large introduction always led to a greater establishment probability, while Grevstad (1999) found several small introductions to be more successful. Further, Wittmann et al. (2014) show that population establishment occurs faster with fewer larger introductions in populations with growth rates below one or with growth rates that fluctuate above and below one. Wittmann et al. (2014) also show that population establishment is faster with several smaller introductions when population growth rate is consistently greater than one. However, our expected mean population growth rate was relatively low (*R* = 1.132; Supplementary Table 2) and we still found that several small introductions had the greatest colonization success. Also, several small introductions performed best in both the fluctuating environment (population growth fluctuated above and below one; Supplementary Table 1) and in the stable environment (population growth consistently greater than one; Supplementary Table 1) (Figure 1, Figure 2). Thus, our results contradict the notion that net reproductive rate is the key control on how introduction regime affects colonization success, which merits further investigation.

We observed striking differences in the measures of colonization success between the microcosm experiment and the demographic simulations. We found that establishment probability increased with the number of introduction events in both the experiment and the simulations, but that all experiment establishment probabilities equaled or exceeded expectations from simulations. In the experiment, mean population size in stable environments was greater than in fluctuating environments, and there was an interaction between environment and introduction regime whereby the mean population size was increasingly greater in stable compared to fluctuating environments as the number of introductions increased. Furthermore, populations grew larger by generation 7 in the experiment than in the simulations.

Differences between the results of the simulations and of the microcosm experiment suggest that the demographic processes captured by the model do not account for all of the biological processes that occurred in the microcosm. Recent work shows that adaptation to the novel, harsh environment from standing variation is possible in this species (Hufbauer et al. 2015), and likely explains the greater establishment probability and population sizes in the microcosm compared to expectations derived from demographic simulations which do not include adaptation.

The rescue effect of immigration can act demographically by increasing the size of populations (Hufbauer et al. 2015). Certainly, demographic rescue played a critical role for the 104 populations that went extinct temporarily until another introduction event revived them. Those temporary extinctions had lasting effects on colonization success. Colonization success declined for populations that experienced a temporary extinction, and the mean population size significantly increased if more colonists contributed to the population after a temporary extinction. These results reflect the overarching importance of total propagule pressure regardless of introduction regime.

The rescue effect can also act genetically by increasing the fitness of populations (Frankham 2015, Hufbauer et al. 2015, Whiteley et al. 2015). The experimental immigrants all came from the same source population, so it is unlikely that gene flow from migrants united previously separated alleles into high-fitness genotypes (Novak 2007). Thus, a more likely mechanism by which immigration increased mean population size beyond expectations was by relieving inbreeding depression or counteracting drift-induced allele loss. Small populations are more prone to experiencing increased homozygosity and inbreeding depression, which can reduce population growth rates and increase extinction risk (McCauley and Wade 1981, O’Grady et al. 2006, Szűcs et al. 2017). Even small amounts of gene flow can alleviate these effects, so the additional small introductions of mated individuals from the external source population were well-suited to bring about longer-term relief (Slatkin 1985, Hufbauer et al. 2015). However, Cassey et al. (2014) found that simulated inbreeding depression was especially detrimental for several, small introductions through time, so other mechanisms are likely at play.

One such mechanism may be adaptation of the incipient population, which is affected by sustained immigration. Introductions to a harsh, novel habitat can result in adaptive evolution with the right amount of gene flow if additional immigrants to a declining population prevent extinction long enough to allow for adaptation to occur (Holt and Gomulkiewicz 1997). However, too much gene flow can lead to genetic swamping whereby the homogenizing effects of gene flow overpowers ongoing local adaptation (Lenormand 2002). This may have been the case for populations in the 10×2 introduction regime, which had the highest average migration rate, lowest establishment probability, and lowest mean population size. Alternatively, negative density dependence may have reduced population fitness when migration rates were high, reducing population growth rates and hampering the spread of adaptive alleles (Holt and Gomulkiewicz 1997). Though not significant, the lower establishment probability and mean population size in the 10×2 introduction regime warrants further work to assess whether they exemplify the yet-unseen scenario described by Blackburn et al. (2015) in which maladaptive gene flow from multiple introductions hampers population establishment in a novel range. More broadly, further study is necessary to evaluate how immigration affected adaptation in this system, if at all (Gomulkiewicz and Holt 1995, Boulding and Hay 2001).

## CONCLUSION

Our experimental results suggest that several, small introductions through time lead to greater colonization success in a novel habitat, and that introductions into a stable recipient environment lead to larger population sizes, but not greater establishment probability. Further, introductions to a stable recipient environment are especially beneficial to populations established with more introduction events. These results defied our expectations derived from parallel simulations of a model that included demographic processes but not evolutionary ones, so we suspect a genetic mechanism might be at work. Genetic mechanisms are rarely incorporated when simulating the effect of introduction regime on colonization (but see Cassey et al. 2014), and our multi-generation microcosm is unique in bringing evolutionary processes to bear on parsing two key components of propagule pressure in an experimental setting.

For invasions, our results highlight the importance of preventing further introductions to the same location, even for established species. For conservation and biological control, our results suggest that emphasis should be placed on increasing the number of introductions or re-introductions to a location, rather than increasing the size of those events if Allee effects are weak. Sustained introduction efforts should also bring about concomitant benefits in the form of longer-term monitoring, increased data collection, and more opportunities for experimentation and adaptive management (Lockwood et al. 2005, Godefroid et al. 2011).

## ACKNOWLEDGMENTS

We thank Charlotte Hoover, Kaitlyn Adkins, Mary Catherine Cochran, Don Waters, Stacy Endriss, Marianna Szűcs, and Jenny Rickford for help with data collection. Funding was provided by NSF Graduate Research Fellowship Grants #DGE-1321845 Amend. 3 (to MJK) and #DGE-1106400 (to MFO), NSF Grant DEB-0949619 (to RAH), and NSF Grant DEB-0949595 (to BAM). RAH also acknowledges the support of the USDA via the Colorado Experiment Station. We thank the lab groups of Ruth Hufbauer, Paul Ode, Andrew Norton, Andrew Latimer, Truman Young, and Jenny Gremer for comments on the project and manuscript.

